# Local and Spatial Processes Shape the Collapse and Recovery of Mutualistic Networks

**DOI:** 10.1101/2025.11.25.690369

**Authors:** Subhendu Bhandary, Jordi Bascompte

## Abstract

Ecosystems can undergo abrupt critical transitions even when environmental change is gradual, resulting in hysteresis, where recovery requires conditions far more favorable than those that triggered collapse. While previous research has mainly focused on local network dynamics, the role of spatial heterogeneity and dispersal in mutualistic ecosystem resilience remains less explored. This study examines how spatial structures—grid, random, small-world, and scale-free networks—interact with dispersal rates to influence ecosystem recovery and mutualistic network persistence. We find that with low dispersal, restoration cost is similar across spatial structures, but at intermediate dispersal rates, scale-free networks show faster recovery and smaller restoration costs. At high dispersal rates, increased connectivity initially reduces restoration cost; however, over time, homogenization weakens spatial heterogeneity, causing restoration cost to increase. Importantly, increasing nestedness can delay collapse but also extends the recovery distance, making ecosystems harder to restore. By adjusting dispersal rates and transitioning from homogeneous to heterogeneous spatial structures, we can decrease restoration cost and improve ecosystem stability, offering key insights for ecological management strategies.

## Introduction

Ecological systems are increasingly destabilized by accelerating global change, driven by habitat fragmentation (Haddad et al., 2015; Wilcox and Murphy, 1985), climate warming (Walther et al., 2002), nitrogen deposition, and widespread deforestation (Laurance et al., 2014). These stressors can push ecosystems past critical thresholds, resulting in sudden and often irreversible regime shifts. Such critical transitions, also referred to as tipping points, are typically driven by nonlinear feedback mechanisms, where the ecosystem abruptly collapses once a bifurcation threshold (environmental stressor) is crossed (Dakos and Bascompte, 2014; Scheffer et al., 2001). Recovery from such a state is often slow and may not occur under the same environmental conditions that caused the collapse. Instead, the system typically requires substantially more favorable conditions to return to its original state—a dynamic known as hysteresis, where collapse and recovery follow different pathways (Kéfi et al., 2014; Scheffer and Carpenter, 2003). Understanding the conditions that govern collapse and recovery, and identifying the factors that influence the cost of restoration, has become central to the study of ecological resilience (Biggs et al., 2018; Suding et al., 2004).

While tipping points have been explored in a variety of systems, including shallow lakes (Carpenter et al., 1999; van de Leemput et al., 2015), coral reefs (McCook, 1999), and savannas (Walker, 1995), mutualistic communities have only recently been examined through this lens. These systems, such as plant–pollinator networks, are characterized by positive feedback loops, where species benefit from the presence of their interaction partners (Bascompte and Jordano, 2007; Bastolla et al., 2009). Such feedbacks promote coexistence under stable conditions but can also increase vulnerability to abrupt collapse when environmental stress intensifies (Aparicio et al., 2021; Bascompte and Scheffer, 2023; Dakos and Bascompte, 2014; Lever et al., 2014). Globally, pollinator populations are declining due to a combination of stressors, including pesticide use, habitat loss, and emerging pathogens (Henry et al., 2012; Potts et al., 2010; Whitehorn et al., 2012). These pressures elevate species mortality and may push mutualistic systems toward critical thresholds or collapse. Predicting community responses to such stress is difficult, as it depends not only on species traits but also on the structure, strength, and organization of their interactions (Bascompte et al., 2006; May, 1972; McCann, 2000). Such findings emphasize the need to study how specific environmental stressors impact the organization of species interactions, and how resulting changes in network structure affect community stability and recovery after disturbance. Network properties such as nestedness and connectance have been shown to influence the robustness and stability of mutualistic communities by determining how interaction strength is distributed across species (Bascompte and Jordano, 2013; Thébault and Fontaine, 2010). However, most studies on tipping points in mutualistic systems have focused on non-spatial, isolated networks, overlooking the spatial complexity inherent in real-world ecosystems (Aparicio et al., 2021; Dakos and Bascompte, 2014; Lever et al., 2014; Panahi et al., 2023).

In nature, species and their interactions are embedded within spatially-structured landscapes, where local communities occupy discrete habitat patches connected by dispersal. These landscapes form metacommunities, where local and regional processes jointly determine biodiversity dynamics (Leibold et al., 2004; Loreau et al., 2003). Dispersal can buffer systems against collapse by enabling recolonization and rescue effects, yet excessive connectivity can synchronize population dynamics across patches, reducing spatial insurance and amplifying the risk of system-wide failure (Urban and Keitt, 2001; Wang and Loreau, 2014). The configuration of the spatial network—whether regular, random, or scale-free—can strongly influence how disturbances spread or are contained within the system (Gilarranz et al., 2015; Holland and Hastings, 2008). Recent work has shown that the spatial organization of landscapes can fundamentally alter ecological tipping points, with heterogeneous structures often delaying collapse and enhancing recovery by promoting localized buffering and spatial insurance (Saade et al., 2023). Despite the increasing recognition of spatial processes in ecological resilience, how local interaction structures (e.g., nestedness) interact with global spatial organization and dispersal to govern collapse, recovery, and hysteresis remains poorly understood. In particular, the degree to which spatial heterogeneity can reduce restoration costs and mediate resilience in mutualistic metacommunities is largely unexplored, representing a key gap this study aims to address.

Recent advances in complex systems theory have shown that multi-layer or network-of-networks frameworks can offer valuable insights into the behavior of interdependent systems (Buldyrev et al., 2010; Kivelä et al., 2014). In such a frameworks, different network layers—whether ecological, infrastructural, or social—are coupled through dynamic processes like information flow or dispersal (Boccaletti et al., 2014; Gao et al., 2012). In ecological contexts, dispersal effectively links local communities across space, forming a multilayer structure where the collapse or recovery of one patch can cascade through the system (Baruah, 2022; Fronhofer et al., 2023). Studies on interdependent systems suggest that structural heterogeneity and the formation of new connections across layers can delay tipping points and enhance system-wide robustness (Altermatt and Fronhofer, 2018; Carrara et al., 2012). By analogy, spatial heterogeneity and moderate dispersal in ecological metacommunities may buffer local collapses and lower restoration costs by redistributing resilience across scales.

In this study, we investigate how the interaction between local mutualistic structure, spatial network topology, and dispersal rate governs collapse, recovery, and hysteresis in mutualistic metacommunities. We model a system in which each habitat patch hosts an plant–pollinator network embedded within spatial configurations of varying complexity (grid, random, small-world, and scale-free). By combining synthetic simulations with 115 empirical networks from the Web of Life database (Fortuna et al., 2014), we quantify how environmental stress influences the cost of restoration and the conditions for recovery. We further evaluate the relative contributions of local and spatial structures to resilience across dispersal regimes using quantitative model comparisons. Our results reveal that spatial heterogeneity fundamentally alters the role of local architecture, with scale-free landscapes under intermediate dispersal most effectively minimizing hysteresis. Together, these insights highlight the need to integrate local interaction structure with spatial network organization to better understand and manage the resilience of mutualistic systems in changing environments.

## Methods

### Spatial metacommunity framework

We modeled a spatially-explicit metacommunity in which each habitat patch harbors a mutualistic community composed of plants and pollinators. These local communities are embedded in a spatial network that governs the dispersal of individuals between patches. Each patch contains a mutualistic interaction network, allowing us to isolate the effects of spatial structure and dispersal. We systematically varied the underlying spatial configuration by generating networks with four distinct topologies: regular (grid-like), small-world, random, and scale-free. This progression introduces increasing heterogeneity in the degree distribution of patches. All spatial structure comprised *N* = 101 patches connected by 202 links, maintaining a constant average degree of 4.

### Local mutualistic dynamics

Within each patch, species dynamics were governed by a set of ordinary differential equations describing population changes for *S_P_* plant and *S_A_* pollinator species (Lever et al., 2014). The abundance of species *i* in patch *l* is denoted by 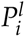 for plants and 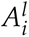 for pollinators. The deterministic dynamics are given by:

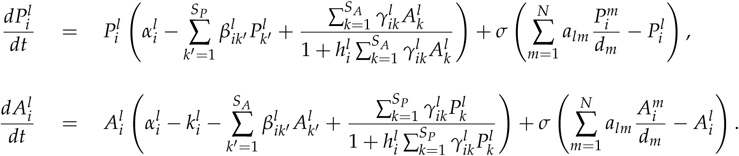

Here, *α_i_* represents the intrinsic growth rate (*α_i_* = 0.15), and *k_i_* denotes an externally imposed mortality affecting only pollinators. Intraspecific and interspecific competition coefficients are set to *β_ii_* = 1 and *β_ij_* = 0.05, respectively. Mutualistic benefits saturate as partner abundance increases, modeled through a type II functional response with handling time 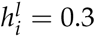.

The mutualistic strength *γ_ik_* is modulated by the species’ connectivity as:

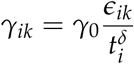

where *ɛ_ik_* represents the entries of the plant pollinator interaction matrix (obtained from 115 empirical networks), *γ*_0_ sets the baseline mutualistic strength (*γ*_0_ = 1), *t_i_* is the degree of species *i*, and *δ* = 0.5 introduces a trade-off between specialization and per-interaction benefit. For simplicity, the main simulations assumed identical parameter values across species and patches; however, we verified that the qualitative patterns are robust to variation in mutualistic strength (Figure S2), local-network heterogeneity (Figure S3), stochastic perturbations (Figure S4), and heterogeneous species-level parameters (Figure S5).

### Dispersal within spatial networks

Dispersal is modeled as a random-walk diffusion process, affecting both plant and pollinator species. The network of spatial connections is encoded by an adjacency matrix *a_lm_*, where *a_lm_* = 1 indicates a direct link between patches *l* and *m*, and *d_m_* is the degree of patch *m*. The net movement of species *i* into patch *l* is given by:

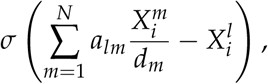

where 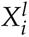 corresponds to either 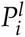 or 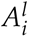, and *σ* is the dispersal rate, varied between 0 (no dispersal) and 1 (high dispersal). This formulation assumes uniform diffusion across neighboring patches, such that individuals spread equally from a given patch to all its neighbors. In regular networks, the diffusion term cancels out under uniform abundance, whereas in heterogeneous networks, local differences in connectivity generate directional flow, influencing spatial persistence and recovery dynamics.

### Simulation protocol and hysteresis quantification

To investigate how the system responds to environmental stress, we incrementally increased the mortality parameter *k_i_* in steps of 0.01, integrating the system numerically using a fourth-order Runge–Kutta solver until a steady state was reached. Extinction was defined as the point where all pollinator abundances dropped below a threshold of 0.01. Following collapse, we reversed the process by gradually decreasing *k_i_* to determine the conditions under which the system could recover. Recovery was defined as the point where at least one pollinator species exceeded the same abundance threshold. The distance between the collapse and recovery points defines the hysteresis width, which we interpret as a measure of the restoration cost. Each simulation involved solving a system of *N* × (*S_P_* + *S_A_*) coupled differential equations.

### Empirical networks

To examine the generality of our findings, we analyzed 115 empirical plant–pollinator networks obtained from the Web of Life database (www.web-of-life.es). We restricted our dataset to networks containing no more than 100 species in total (plants plus pollinators) to maintain computational feasibility while preserving a broad spectrum of structural diversity. These empirical systems encompass a variety of interaction patterns—from highly specialized to strongly generalized—providing a representative gradient of local network organization through which to assess spatial and dispersal effects on system resilience.

We quantified the internal organization of each empirical network using nestedness, a key descriptor of mutualistic network that captures the tendency of specialist species to interact with subsets of generalist partners. To quantify this pattern, we used the metric proposed by (Fortuna et al., 2019), which is mathematically equivalent to the NODF measure (Almeida-Neto et al., 2008) but avoids penalizing species that share the same number of interaction partners. For each pair of plant or pollinator species, we calculated the proportion of shared interaction partners relative to the smaller of their degrees and then averaged these values across all pairs to obtain the observed nestedness of each network. To allow comparisons among networks differing in size, connectance, or sampling intensity, we standardized nestedness using a null model–based *z*-score. Following the probabilistic null model (Bascompte et al., 2003) randomized matrices were generated by drawing each cell independently with probability *π_ij_* = (*p_i_* + *q_j_*)/2 where *p_i_* and *q_j_* denote the empirical row and column marginal frequencies of plant and pollinator respectively. This null model preserves the expected generalization level of both plants and pollinators, while allowing variation around the empirical structure. For each empirical network, we generated 100 randomized matrices computed their mean (*µ*) and standard deviation (*σ*) of nestedness and computed the standardized nestedness value as 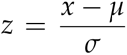, where *x* denotes the observed nestedness of the empirical network. This normalization allows direct comparison of nestedness across systems differing in size, connectance, and sampling intensity. The resulting nestedness *z*-score thus reflects how strongly a network deviates from random expectations—positive values indicating more nested organization than expected by chance.

To evaluate the influence of local structural variability on resilience under a fixed spatial configuration, we systematically embedded each one of the 115 empirical networks—each characterized by a distinct nestedness *z*-score—within the same spatial topologies. This approach allowed us to isolate how differences in local mutualistic architecture modify system behavior when the spatial framework is held constant. In practice, for a given spatial structure (e.g., grid, random, small-world, or scale-free), each empirical network served as the local interaction matrix for all patches, while dispersal dynamics and global connectivity patterns remained unchanged. This design ensured that any observed variation in collapse thresholds, recovery trajectories, or restoration costs arose specifically from differences in local network organization, rather than from alterations in the spatial structure or dispersal regime. By repeating this process across multiple spatial configurations, we systematically disentangled the relative contributions of local architecture and spatial topology to metacommunity resilience.

## Results

### Dispersal Rates and Spatial Structures in Collapse and Recovery Dynamics

We observed that the impact of dispersal on collapse and recovery dynamics varied across spatial structures (Figure 2). In the absence of dispersal, collapse points were nearly identical across all networks, as spatial configuration could not influence dynamics without connectivity among patches. At intermediate dispersal rates, scale-free networks exhibited a delayed collapse and faster recovery compared with grid and random networks, reflecting the stabilizing role of heterogeneous connectivity (Figure 2, S1). At high dispersal, scale-free systems collapsed and recovered earlier, indicating that strong coupling accelerates both loss and recovery through homogenization. The hysteresis width, representing restoration cost, declined with increasing dispersal but rose again beyond a critical threshold, suggesting diminishing stabilizing effects at very high dispersal. These results highlight intermediate dispersal as the regime that most effectively balances stability and recovery potential across spatial configurations.

**Figure 1:**
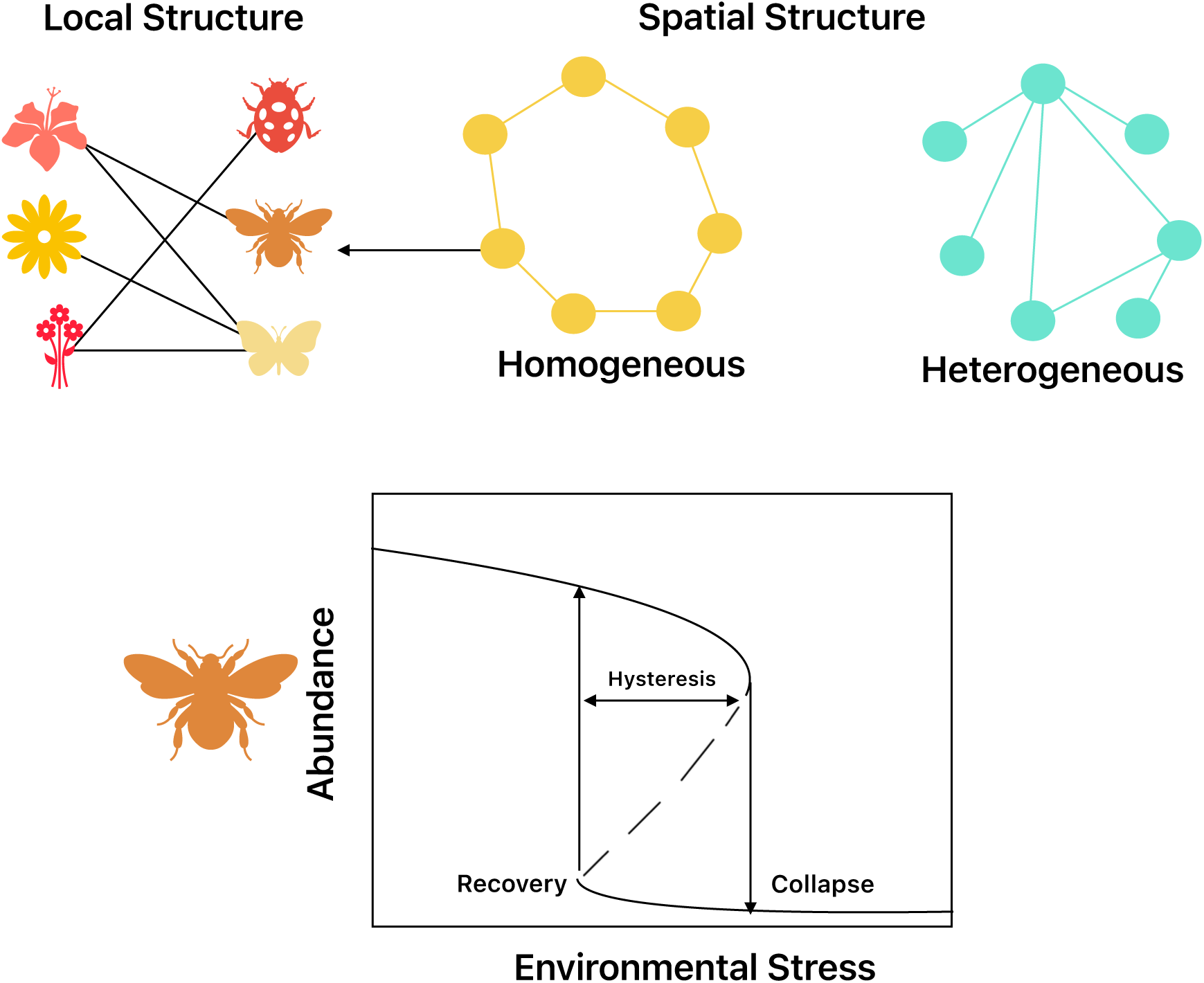
Schematic of local and spatial dynamics in mutualistic metacommunities. Local mutualistic networks of plants and pollinators are embedded within spatial networks that range from homogeneous (grid) to heterogeneous (scale-free) structures. The lower panel illustrates how increasing environmental stress drives a collapse in species abundance, followed by recovery as stress is reduced, forming a hysteresis loop. The distance between collapse and recovery thresholds represents the restoration cost, highlighting how the interplay between local interactions and spatial connectivity governs the recovery of mutualistic systems.

**Figure 2:**
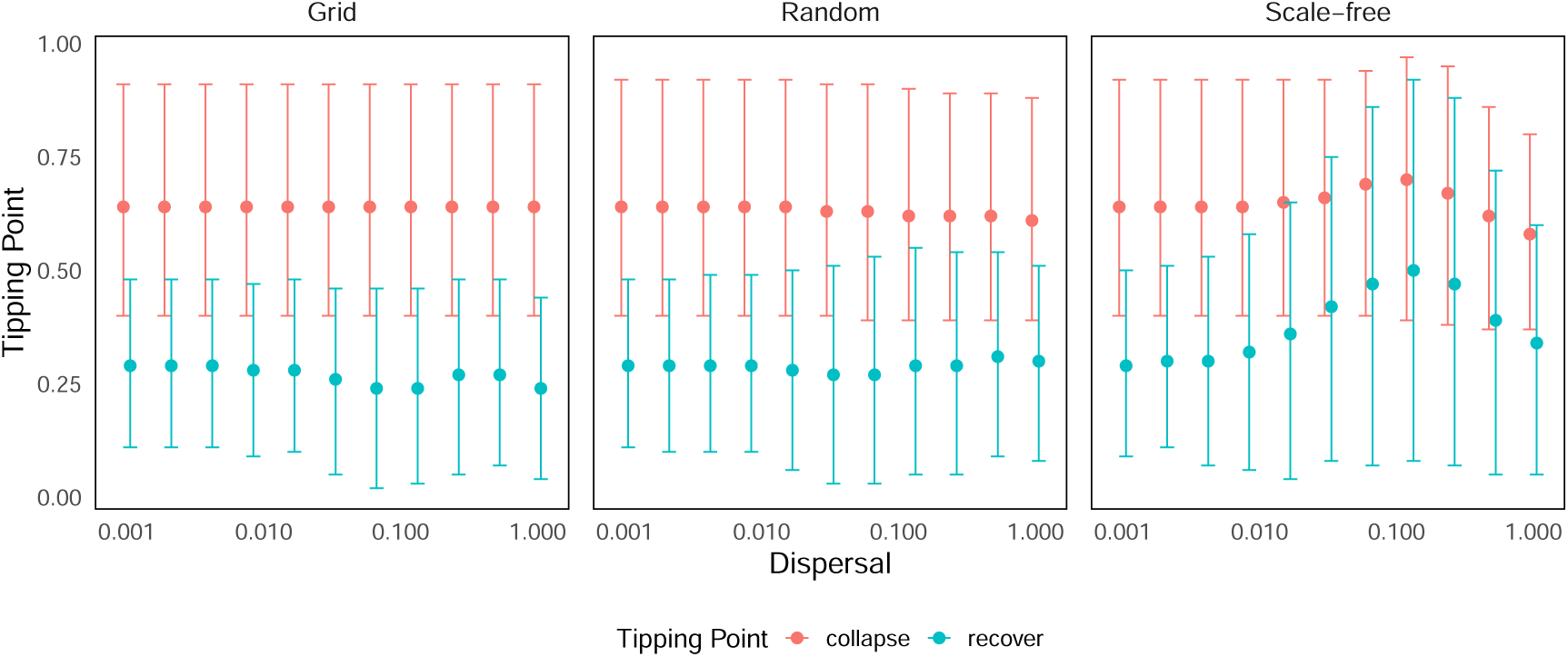
Collapse and recovery thresholds across spatial and dispersal regimes. Each subplot presents results from 115 mutualistic network realizations across three spatial configurations (grid, random, and scale-free) and varying dispersal rates. Filled circles indicate the mean collapse and recovery points, while error bars represent the 95% confidence intervals. The figure highlights that at intermediate dispersal, scale-free networks experience a delayed collapse and a faster recovery relative to grid and random structures.

Building on the observed collapse and recovery dynamics, we next examined how local network architecture—specifically nestedness—modulates restoration cost (hysteresis width) across spatial structures and dispersal regimes (Figure 3). The first row of Figure 3 shows how restoration cost varies with nestedness for grid, random, and scale-free networks under low (*σ* ≈ 0), intermediate (*σ* = 0.125), and high dispersal (*σ* = 1), while the second row presents the corresponding regression slopes that quantify the direction and strength of these relationships.

**Figure 3:**
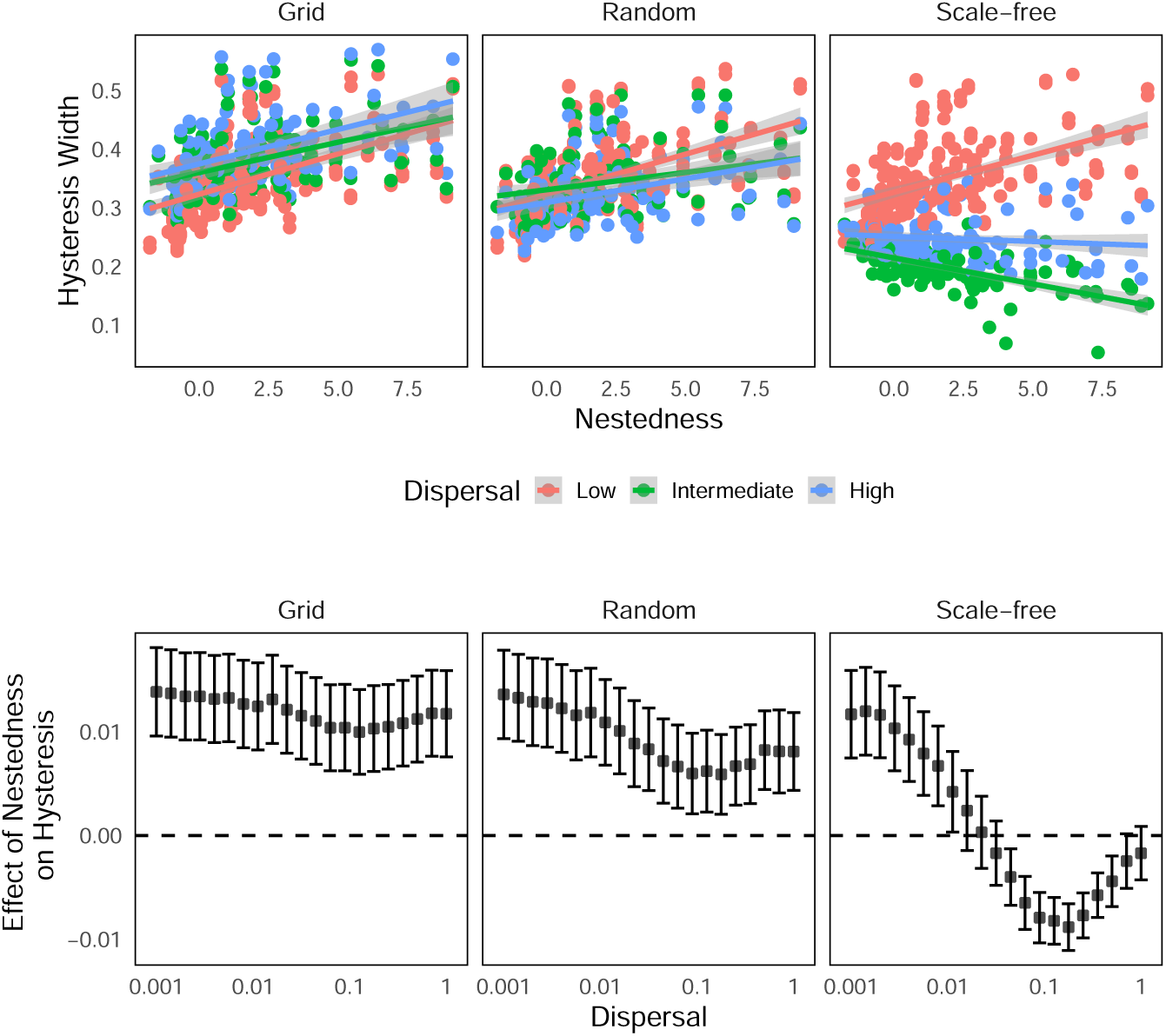
Variation in restoration cost with nestedness across spatial structures and dispersal regimes. The first row shows how restoration cost (measured as hysteresis width) varies with nestedness across three spatial structures—grid, random, and scale-free—under low (red), intermediate (green), and high (blue) dispersal rates. Linear model fits illustrate the direction and strength of these relationships. The second row depicts the estimated slopes from these models, highlighting whether nestedness increases or decreases restoration cost; dashed lines mark zero to distinguish positive and negative effects. Grid and random networks generally exhibit a positive association between nestedness and restoration cost, whereas scale-free networks show a dispersal-dependent shift—nestedness lowers restoration cost at intermediate dispersal, indicating enhanced recovery efficiency under heterogeneous spatial connectivity.

In grid and random networks, restoration cost increased consistently with nestedness across all dispersal levels, reflecting a persistent positive slope. In grids, this relationship remained equally strong regardless of dispersal, whereas in random networks the positive slope weakened at intermediate dispersal before strengthening again at high dispersal. In contrast, scale-free networks showed a clear dispersal-dependent reversal: the slope was positive at low dispersal, became negative at intermediate levels—indicating that moderate movement through heterogeneous connectivity reduced restoration cost—and approached zero at high dispersal as spatial mixing homogenized the system. Among all structures, grid networks exhibited the steepest positive slopes, followed by random and then scale-free networks. The lowest restoration cost occurred in highly nested scale-free systems at intermediate dispersal, marking the regime where spatial heterogeneity and moderate connectivity most effectively enhance recovery efficiency.

### Local and Spatial Contributions to Restoration Cost

To evaluate how local and spatial structures jointly influence restoration cost, we mapped hysteresis width against nestedness (local property) and spatial configuration across dispersal regimes (Figure 4). At low dispersal, variation in restoration cost was mainly associated with nestedness, indicating that local network architecture dominated resilience when movement between patches was limited. With increasing dispersal, patterns progressively shifted toward the spatial axis, showing that spatial connectivity—particularly in scale-free landscapes—became the principal driver of reduced restoration cost. This transition marked a threshold where spatial organization began to govern recovery dynamics. At high dispersal, the effects of both local and spatial structure converged, reflecting the homogenizing influence of strong connectivity. Overall, intermediate dispersal emerged as the regime where spatial heterogeneity most effectively reduced restoration cost and enhanced system resilience.

**Figure 4:**
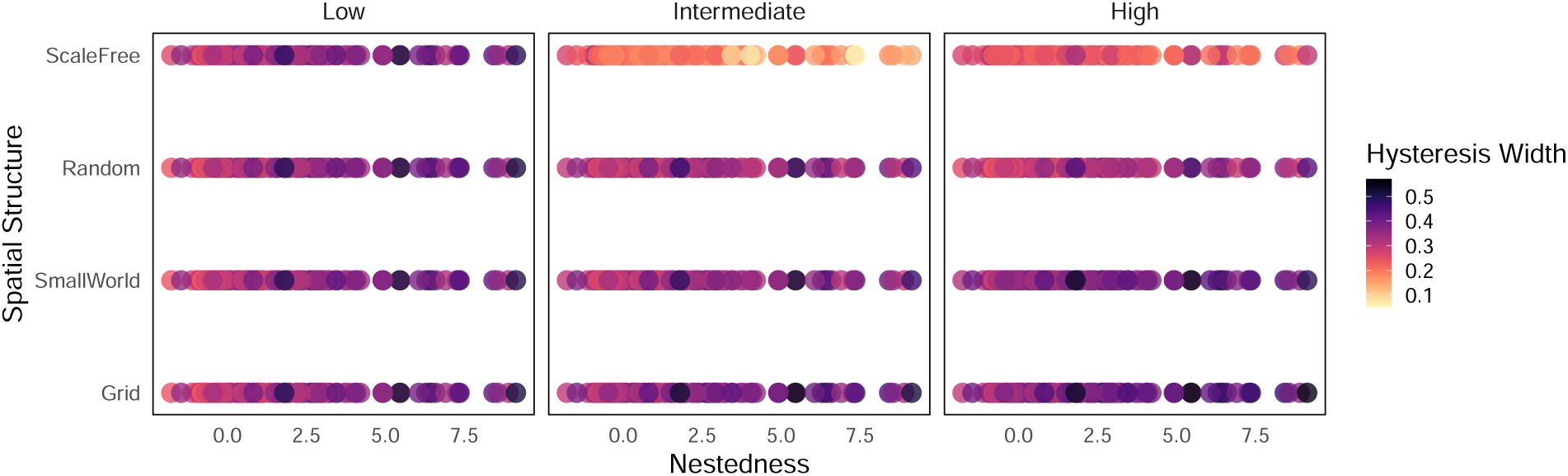
Heatmap illustrating the combined effects of local and spatial structures on restoration cost across dispersal regimes. Restoration cost (hysteresis width) is shown as a function of local network nestedness (x-axis) and spatial structure type (grid, random, scale-free; y-axis) for varying dispersal rates. At low dispersal, variation in restoration cost is largely governed by local structure, reflecting patch-level control of resilience. With increasing dispersal, restoration cost patterns shift toward the spatial axis, indicating that network connectivity—particularly in scale-free structures—enhances recovery and lowers restoration costs. At high dispersal, the influence of both axes converges, consistent with landscape homogenization. Overall, the figure highlights that intermediate dispersal optimally balances local and spatial processes to enhance ecosystem resilience.

We quantitatively evaluated how local and spatial network structures shape restoration cost (measured as hysteresis width) across varying dispersal rates. Using generalized linear models (GLMs) and comparing their performance with Akaike Information Criterion (AIC), we found that models combining both local (nestedness) and spatial (network topology) predictors consistently outperformed single-factor models, underscoring the interdependence of local and global processes (Figure 5). At low dispersal rates, local structural properties exerted stronger positive effects on restoration cost, indicating that limited connectivity amplifies local feedbacks and delays recovery. In contrast, at intermediate dispersal rates, spatial structure became the dominant driver, where heterogeneous connectivity—particularly in scale-free networks—significantly reduced restoration cost. At high dispersal, both effects converged, reflecting homogenization across the landscape that diminished the distinct influence of either structure. Overall, these results demonstrate that local architecture governs resilience under isolation, whereas spatial heterogeneity mitigates restoration cost under moderate dispersal, promoting efficient recovery at the metacommunity scale. Statistical comparisons confirmed that models integrating both local and spatial predictors best explained restoration cost, emphasizing the synergistic-not additive-nature of their effects.

**Figure 5:**
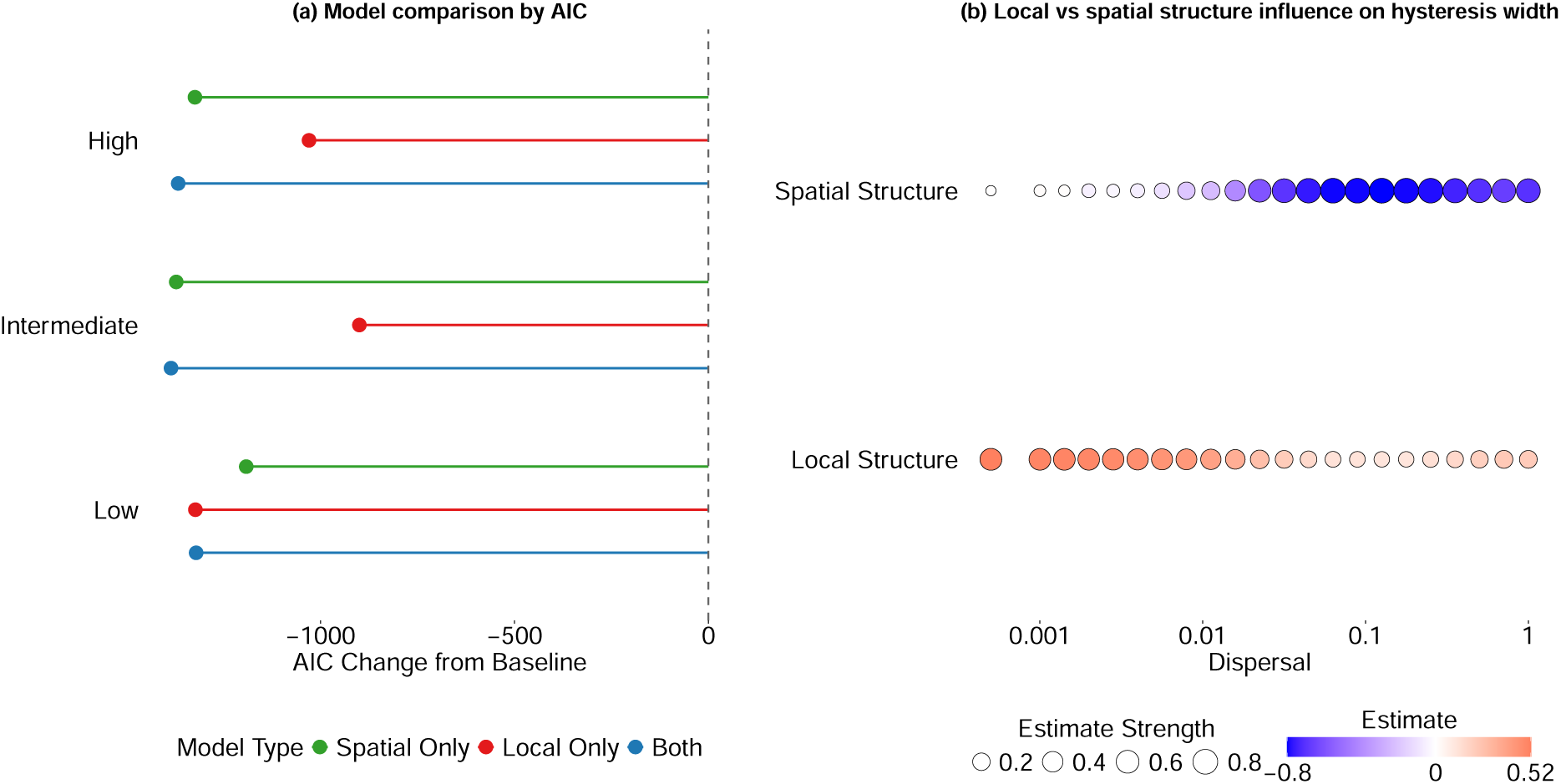
Quantitative evaluation of local and spatial contributions to restoration cost across dispersal regimes. (a) Model comparison using Akaike Information Criterion (AIC) across low, intermediate, and high dispersal rates. Models incorporating both local (nestedness) and spatial (network topology) predictors consistently yielded lower AIC values than single-factor models, indicating that restoration cost (hysteresis width) is jointly governed by local and spatial factors. (b) Standardized regression coefficients (*β*) for local and spatial predictors across dispersal regimes. Point color represents the direction and magnitude of effects, while size denotes effect strength. At low dispersal, local structure strongly increases restoration cost, reflecting dominant local feedbacks under limited connectivity. At intermediate dispersal, spatial structure becomes the primary driver, where heterogeneous connectivity markedly reduces restoration cost. At high dispersal, both effects converge, reflecting landscape homogenization. Overall, while local structure amplifies restoration cost in isolated conditions, spatial heterogeneity mitigates it under moderate dispersal, promoting more efficient recovery at the landscape scale.

## Discussion

Our work shows that resilience in mutualistic metacommunities arises from the dynamic interaction between local mutualistic structure and the architecture of spatial connectivity. By combining 115 empirical plant–pollinator networks with multiple spatial configurations and dispersal regimes, we reveal how these two scales jointly control collapse, recovery, and the cost of restoration. Dispersal mediates a transition in control—from recovery governed by local interactions at low connectivity to spatially-coordinated stabilization at intermediate dispersal, where heterogeneous, scale-free landscapes most effectively reduce hysteresis. Unlike previous studies that analyzed local mutualistic networks in isolation, our framework integrates spatial heterogeneity and dispersal processes, demonstrating that stability emerges not from either local or spatial structure alone, but from their coupling. This integration identifies the dispersal regime where spatial heterogeneity most strongly promotes recovery and minimizes restoration cost.

Abrupt transitions in ecosystems typically emerge from reinforcing feedbacks between species interactions and environmental stress (Dakos and Bascompte, 2014; May, 1972; Scheffer et al., 2001). Although these dynamics are well documented in isolated communities, real landscapes consist of spatially-distributed patches linked through dispersal (Leibold et al., 2004; Loreau et al., 2003). Spatial connectivity can either buffer collapse by facilitating recolonization or propagate disturbances that synchronize declines (Dakos et al., 2019; Gilarranz et al., 2015). Our results extend this theoretical foundation by quantifying how spatial topology and dispersal together determine restoration cost—a metric that reflects how much effort is required to return a system to its original state after collapse. By explicitly coupling local nestedness with spatial structure, we identify how landscape heterogeneity governs the balance between local persistence and regional synchronization (Holland and Hastings, 2008; Kéfi et al., 2014; Saavedra et al., 2017).

At low dispersal, local architecture dominates system behavior: each patch recovers independently, consistent with earlier theoretical predictions (Lever et al., 2014; Rohr et al., 2014). As dispersal increases, however, spatial links allow recovery in one patch to spread to others, lowering restoration cost through a “spatial insurance effect” (Loreau et al., 2003). This effect was strongest in heterogeneous (scale-free) spatial networks, where hub-like nodes channel recovery potential and restrict the propagation of collapse. Excessive dispersal, however, erased spatial differences, synchronized patch dynamics, and widened hysteresis (Dakos et al., 2019; Wang and Loreau, 2014). This dual behavior highlights a general ecological trade-off: dispersal can promote recovery but, when too strong, amplifies systemic vulnerability—a phenomenon also observed in interdependent infrastructures and food webs (Buldyrev et al., 2010; Gao et al., 2012).

From a restoration perspective, our findings clarify how the influence of local and spatial structure shifts with dispersal. While local properties such as nestedness regulate resilience, their effects depend strongly on spatial organization. Highly nested networks embedded in scale-free landscapes exhibited the lowest restoration costs, revealing that spatial heterogeneity can compensate for the recovery inefficiency of densely connected local architectures. Crucially, these patterns were not sensitive to simplifying assumptions: the same qualitative behaviour emerged under environmental heterogeneity, stochastic perturbations, variation in local mutualistic structure, and species-level parameter differences drawn from uniform ranges (Figs. S2–S5). Thus, optimizing restoration in mutualistic systems may require not only local rewiring but also targeted modification of spatial connectivity. Moderate dispersal in heterogeneous landscapes can create recovery corridors without inducing full synchronization,—a principle consistent with metacommunity theory and recent work on spatial design for ecosystem resilience (Bascompte et al., 2019; Keitt et al., 1997).

Our framework has limitations that open directions for further research. Although we introduced heterogeneity in species parameters, all patches in our model experienced the same underlying environmental conditions, whereas in real landscapes demographic rates and interaction strengths are shaped by local abiotic factors such as temperature, moisture, nutrient availability, and disturbance history. Such spatial variation in environmental drivers can generate patch-specific collapse and recovery dynamics (Dakos et al., 2019) that our uniform environmental backdrop does not capture. Adaptive behaviors—such as trait evolution, partner switching (Cai et al., 2020; Kaiser-Bunbury et al., 2010; Kondoh, 2003; Valdovinos et al., 2013) were also not considered, yet these can enhance persistence and modify spatial tipping points (Valdovinos et al., 2016). Furthermore, we focused on purely mutualistic layers, while antagonistic links (e.g., herbivory, parasitism) often coexist and may redistribute extinction risks or stabilize certain modules (Glaum and Kessler, 2017; Pilosof et al., 2017). Finally, dispersal was assumed static and uniform; in nature, movement is adaptive and species-specific, potentially feeding back on spatial structure and recovery trajectories. Extending the model to account for these processes would increase ecological realism and improve predictions of collapse and restoration in fragmented landscapes.

Together, these findings establish that spatial structure has a stronger qualitative role than local architecture in determining restoration efficiency (Figure 5(b)). While increasing nestedness can delay collapse, it may also widen hysteresis by slowing recovery once disruption occurs. Spatial heterogeneity can counterbalance this cost by redistributing recovery potential across the network, effectively reducing the restoration burden. This mechanism mirrors observations in fragmented pollination networks, coral reef systems, and stepping-stone habitats, where moderate connectivity enhances recolonization and stabilizes mutualistic interactions (Keitt et al., 1997; Kormann et al., 2016). By identifying when and how landscape structure alters the resilience of mutualistic communities, this study bridges theoretical ecology with conservation practice. It suggests that maintaining spatial heterogeneity—rather than altering local network design alone—may represent a more efficient and cost-effective path to restoring and sustaining ecosystem function in a changing world.

## Data Availability

Codes and data are available in a Github repository (https://github.com/subhendu-math/Local-spatial-effect.git).

## Acknowledgments

Subhendu thank the members of Bascompte Lab for discussions. Funding was provided by SNSF (grant number 310030 197201 to JB).

## Notes

### Competing Interest Statement

The authors have declared no competing interest.

