## Supporting Information for "Local and Spatial Processes Shape the Collapse and Recovery of Mutualistic Networks"

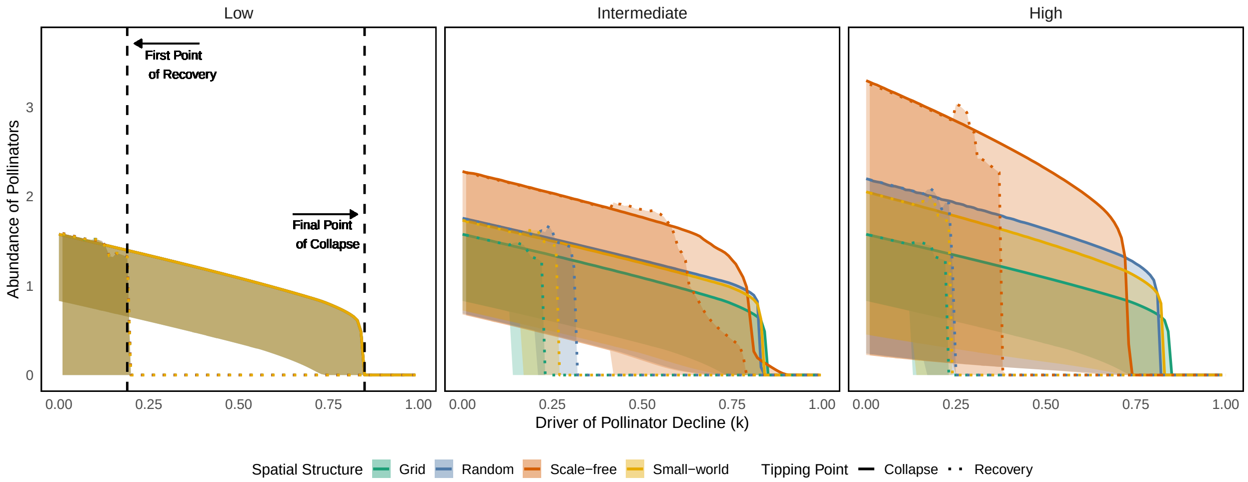


**Figure S1.** Collapse and recovery dynamics across varying pollinator decline rates and dispersal strengths. Each column corresponds to low, intermediate, and high dispersal scenarios. Under low dispersal, collapse and recovery coincide for all spatial structures, shown by dotted reference lines. At intermediate dispersal, the distance between collapse and recovery decreases, with restoration cost lowest for Scale-free, followed by Random, Small-world, and highest for Grid networks. With further increase in dispersal, overall restoration cost rises again, but the structural hierarchy (Scale-free < Random < Small-world < Grid) remains consistent. Color width indicates individual species, solid lines denote final collapse points, and dotted lines mark the first recovery points.

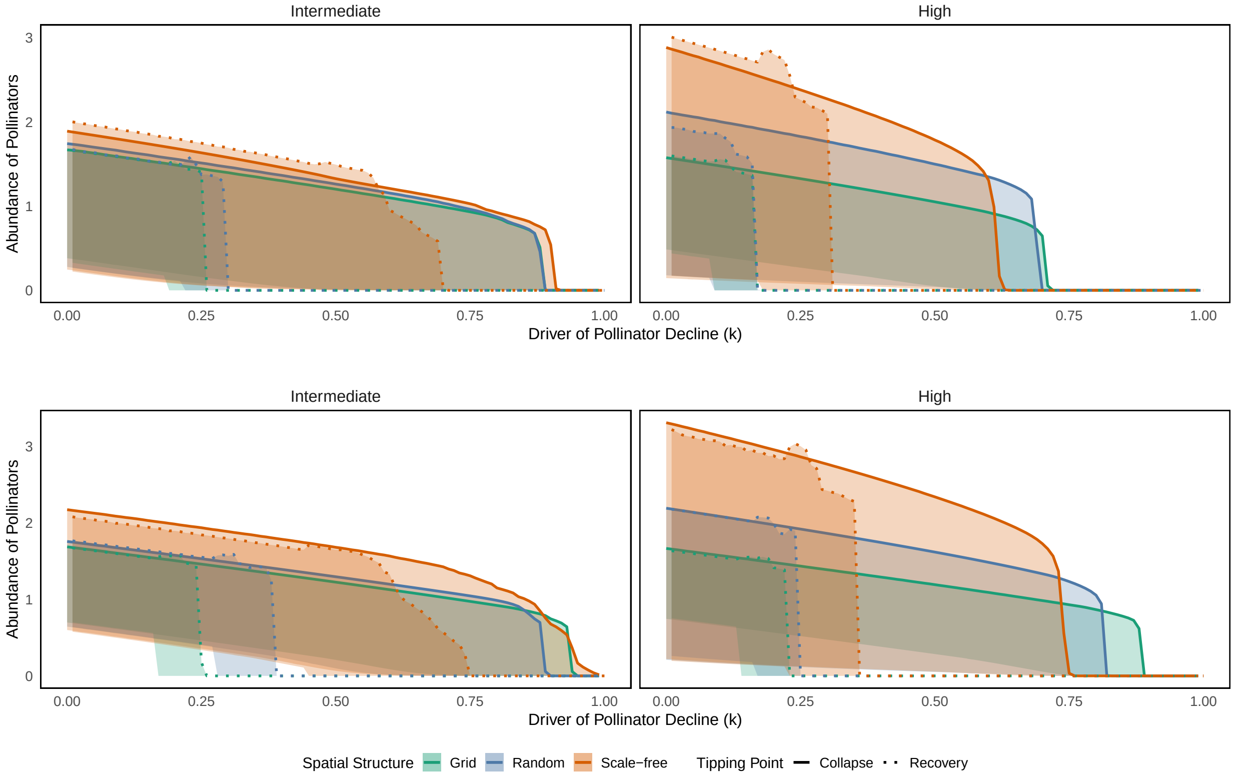


**Figure S2.** Collapse and recovery dynamics under heterogeneous mutualistic strengths across patches and varying dispersal rates. Each subplot shows the pollinators abundance during collapse (solid lines) and recovery (dotted lines) for two spatial structures: Grid (green) and Scale-free (orange). The first row corresponds to randomly assigned mutualistic strengths $\gamma_0 \in [0.4, 1.2]$, and the second row to $\gamma_0 \in [0.8, 1.2]$, across intermediate and high dispersal scenarios. These intervals represent heterogeneous environmental conditions, where each patch is assigned a random mutualistic strength drawn from the specified range. The wider interval, containing lower $\gamma_0$ values, leads to earlier collapse, whereas the narrower interval with higher $\gamma_0$ values delays the critical transitions. In all cases, the scale-free structure consistently exhibits a smaller collapse–recovery gap, indicating reduced restoration cost irrespective of local heterogeneity.

### Robustness of spatial effects under heterogeneous local mutualistic structures

To evaluate whether the influence of spatial structure on collapse and recovery dynamics remains consistent under locally heterogeneous conditions, we generated a set of spatially distributed mutualistic networks with controlled variation in local community composition. Starting from a baseline bipartite interaction matrix, we introduced heterogeneity among local networks by modifying their adjacency matrices while maintaining species richness and connectance. The degree of dissimilarity among patches was quantified using the Bray–Curtis dissimilarity index, which measures changes in the distribution of interactions between species pairs. Specifically, we generated two contrasting levels of heterogeneity: (i) low heterogeneity ($\beta = 0.1$) representing nearly homogeneous local structures, and (ii) high heterogeneity ($\beta = 0.9$) representing highly distinct local networks (see Supplementary Code S3 for network generation details).

Despite substantial variation in local mutualistic architecture, the qualitative patterns of collapse and recovery remained consistent across spatial structures. As shown in Fig. S3, both Grid (green) and Scale-free (orange) spatial networks exhibit a characteristic hysteresis pattern, with solid lines representing collapse trajectories and dotted lines representing recovery. However, the distance between collapse and recovery thresholds—interpreted as restoration cost—remains consistently smaller in the scale-free structure compared to the grid structure, even under high local heterogeneity. This demonstrates that the stabilizing effect of spatial heterogeneity in connectivity (i.e., global structure) persists even when local network organization varies strongly among patches. In other words, the low restoration cost advantage conferred by heterogeneous spatial connectivity is robust to changes in local mutualistic composition.

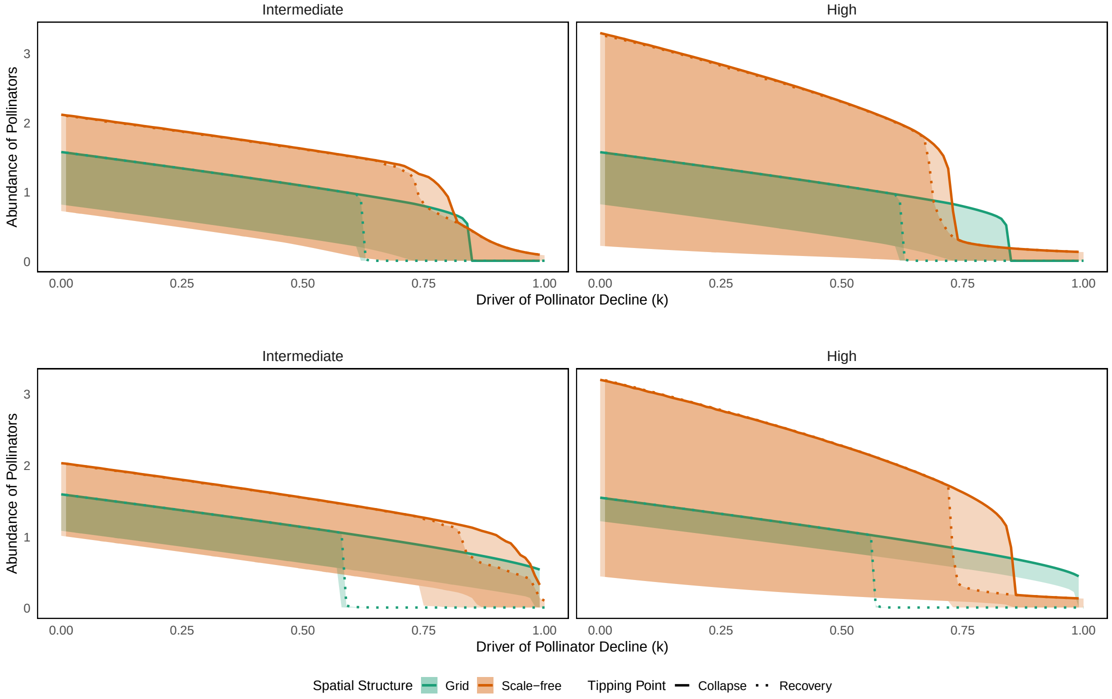


**Figure S3. Effect of local heterogeneity on collapse and recovery across spatial structures.** Each subplot compares the pollinator abundance during collapse (solid lines) and recovery (dotted lines) for two spatial structures: Grid (green) and Scale-free (orange). The first row represents low heterogeneity in local mutualistic networks ($\beta = 0.1$), while the second row represents high heterogeneity ($\beta = 0.9$), both under fixed nestedness. Shaded regions denote variation in species across patches. Despite increasing heterogeneity among local interaction networks, the smaller collapse–recovery gap in the scale-free structure indicates that spatial topology continues to determine system-level restoration cost, confirming the robustness of our results.

### Model robustness under stochastic fluctuations

To assess the robustness of our findings, we extended the deterministic mutualistic metacommunity model by incorporating demographic stochastic noise and diffusive spatial coupling. The noise-augmented dynamics for species $i$ in patch $l$ are given by:

$$\frac{dP_i^l}{dt} = P_i^l\left(\alpha_i^l - \sum_{k'=1}^{S_P}\beta_{ik'}^l P_{k'}^l + \frac{\sum_{k=1}^{S_A}\gamma_{ik}^l A_k^l}{1+h_i^l\sum_{k=1}^{S_A}\gamma_{ik}^l A_k^l}\right) + \sigma\left(\sum_{m=1}^{N} a_{lm}\frac{P_i^m}{d_m} - P_i^l\right) + \sigma_{\text{dem}}\sqrt{P_i^l}\mu_i^l(t),$$

$$\frac{dA_i^l}{dt} = A_i^l\left(\alpha_i^l - k_i^l - \sum_{k'=1}^{S_A}\beta_{ik'}^l A_{k'}^l + \frac{\sum_{k=1}^{S_P}\gamma_{ik}^l P_k^l}{1+h_i^l\sum_{k=1}^{S_P}\gamma_{ik}^l P_k^l}\right) + \sigma\left(\sum_{m=1}^{N} a_{lm}\frac{A_i^m}{d_m} - A_i^l\right) + \sigma_{\text{dem}}\sqrt{A_i^l}\xi_i^l(t).$$

Here $\sigma_{\text{dem}}$ regulates the strength of demographic stochasticity, while $\xi_i^l(t)$ and $\mu_i^l(t)$ are independent white-noise processes with zero mean and unit variance. We use multiplicative noise, scaled by $\sqrt{P_i^l}$ and $\sqrt{A_i^l}$ which links noise intensity to population size and thus captures realistic demographic variability. This formulation captures both environmental heterogeneity and demographic variability, ensuring ecological realism in the system's temporal evolution.

We evaluated two representative regimes: $(i)$ intermediate dispersal with low noise strength ($\sigma_{\text{dem}} = 0.01$), and $(ii)$ high dispersal with stronger noise intensity ($\sigma_{\text{dem}} = 1$). In both regimes, the system exhibited clear collapse and recovery transitions, analogous to the deterministic case. Importantly, the distance between collapse and recovery thresholds remained smaller in scale-free networks compared to grid networks, indicating higher resilience and faster recovery in heterogeneous spatial structures. The qualitative consistency across both noise regimes demonstrates that our results are robust to stochastic perturbations, confirming that the observed differences in tipping behavior between spatial structures are not artifacts of deterministic assumptions.

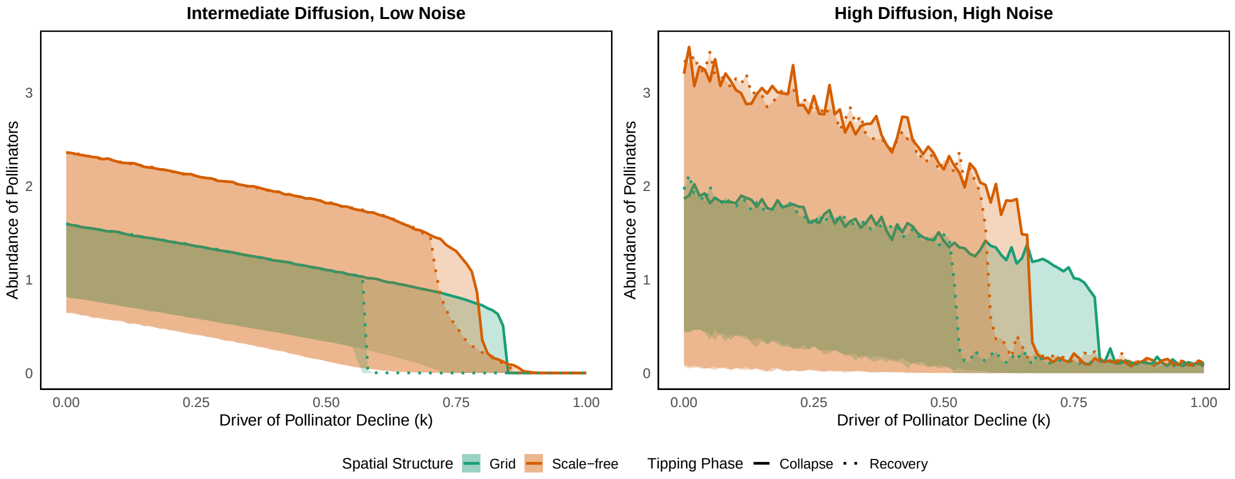


**Figure S4.** Robustness of collapse and recovery dynamics under stochastic fluctuations. Each panel shows pollinator abundance during forward (collapse; solid line) and backward (recovery; dashed line) transitions for two spatial structures—Grid (green) and Scale-free (orange). First column for intermediate dispersal with low noise strength and second column for high dispersal with strong noise demonstrate consistent tipping dynamics across stochastic regimes. Shaded areas represent the range (minimum–maximum) of abundances across all species. The reduced hysteresis gap in scale-free networks compared to grid-like structures indicates greater robustness to perturbations.

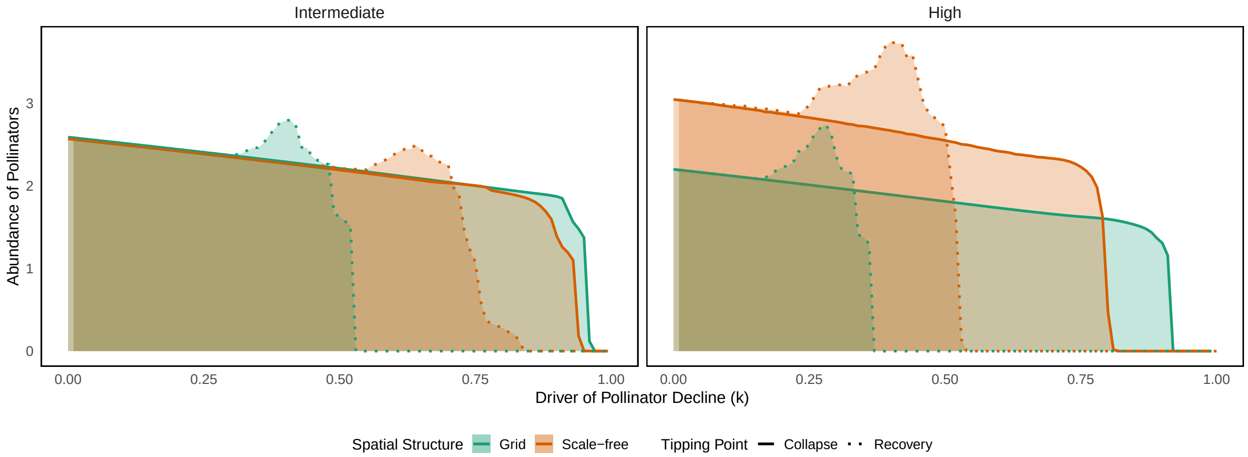


**Figure S5.** Stability of tipping behaviour under heterogeneous species parameters. Forward (solid) and backward (dashed) transitions of pollinator abundance are shown for two spatial configurations: grid (green) and scale–free (orange). The left and right columns correspond to intermediate and high dispersal, respectively. Shaded bands denote the full spread of species abundances. To introduce variation among species, all demographic and interaction parameters were assigned independently from uniform distributions: $\alpha_i^l \sim U(0.05, 0.35)$, $h_i^l \sim U(0.15, 0.30)$, $\beta_{ik}^l \sim U(0.01, 0.05)$, $\beta_{ii}^l \sim U(0.8, 1.1)$, $\gamma_{0,ik}^l \sim U(0.8, 1.2)$. Across this heterogeneous parameter space, scale–free networks maintain a consistently smaller collapse–recovery gap than grid networks, indicating that their resilience advantage persists even when species differ widely in their intrinsic dynamics.
